# A cryogenic, coincident fluorescence, electron and ion beam microscope

**DOI:** 10.1101/2022.09.02.506334

**Authors:** Daan B. Boltje, Jacob P. Hoogenboom, Arjen J. Jakobi, Grant J. Jensen, Caspar T.H. Jonker, Max J. Kaag, Abraham J. Koster, Mart G.F. Last, Cecilia de Agrela Pinto, Jürgen M. Plitzko, Stefan Raunser, Sebastian Tacke, Zhexin Wang, Ernest B. van der Wee, Roger Wepf, Sander den Hoedt

## Abstract

Cryogenic electron tomography (cryo-ET) combined with sub-tomogram averaging, allows *in-situ* visualisation and structure determination of macromolecular complexes at sub-nanometre resolution. Cryogenic focused ion beam (cryo-FIB) micromachining is used to prepare a thin lamella-shaped sample out of a frozen-hydrated cell for cryo-ET imaging, but standard cryo-FIB fabrication is blind to the precise location of the structure or proteins of interest. Fluorescence-guided focused ion beam (FIB) milling at target locations requires multiple sample transfers prone to contamination, and relocation and registration accuracy is often insufficient for 3D targeting. Here, we present *in-situ* fluoresence microscopy-guided FIB fabrication of a frozen-hydrated lamella to solve this problem: we built a coincident 3-beam cryogenic correlative microscope by retrofitting a compact cryogenic microcooler, custom positioning stage, and an inverted widefield fluorescence microscope (FM) on an existing focused ion-beam scanning electron microscope (FIB-SEM). We show FM controlled targeting at every milling step in the lamella fabrication process, validated with transmission electron microscope (TEM) tomogram reconstructions of the target regions. The ability to check the lamella during and after the milling process results in a higher success rate in the fabrication process and will increase the throughput of fabrication for lamellae suitable for high-resolution imaging.

## Introduction

High-resolution 3D reconstructions of biological macromolecules in their near-native cellular environment are necessary to obtain a mechanistic understanding of complex, biological processes at the molecular scale. Cryo-ET allows imaging cellular structures at unprecedented resolution and clarity ***Zimmerli et al. (2021)***; ***Tegunov et al. (2021)***. In cryo-ET, a sample is flash cooled, thinned to the appropriate thickness, and a tilt series of projections is acquired using a TEM ***Koning et al. (2018)***; ***Turk and Baumeister (2020)***. A prerequisite for high-resolution cryo-ET is that the sample is thinner than the inelastic mean-free-path of electrons in vitreous water ice—in practice, this means a sample thickness of approximately 100 to 200 nm ***Vulović et al. (2013)***. To create sufficiently thin sections (lamellae) for high-resolution tomography, the use of a FIB-SEM has proven to be most successful in ablating the excess cellular material surrounding the region of interest ***Chiang et al. (2007)***; ***Villa et al. (2013)***; ***Hylton and Swulius (2021)***. In recent years, various improvements and refinements have been made to the cryo-FIB milling workflow improving throughput, reliability, sample yield, and quality ***Schaffer et al. (2019)***; ***Wolff et al. (2019)***; ***Tacke et al. (2021)***; ***Buckley et al. (2020)***.

Identification of the region of interest (ROI) for milling in the right location is a crucial step. Unfortunately, neither the scanning electron microscope (SEM) nor the FIB provides a contrast mechanism for biomolecular composition, leading to blind milling and possibly inadvertent ablation of the structure of interest. Cryogenic correlative light and electron microscopy (cryo-CLEM) can be employed to overcome this challenge. In this approach, the location of specific objects or cellular compartments is targeted for cryo-FIB milling using specific fluorescent labeling, thus allowing targeted FIB milling.

The first use of cryo-CLEM for fluorescence targeted FIB milling employed mono-sized ferromagnetic polystyrene beads as fiducial markers for FM and FIB-SEM correlation ***Arnold et al. (2016)***. The signal from the iron oxide in the magnetic beads is detectable in the FIB-SEM by the backscattered electron detector and the polystyrene is auto-fluorescent when excited by green light in the FM. FM imaging for target localisation was done in a stand-alone cryogenic spinning disk confocal microscope after which the sample was transferred to the FIB-SEM. Correlation was achieved by 3D coordinate transformation and a correlation accuracy on the order of tens of nanometer was shown. However, the error range remained relatively large, ranging from 0.2 to 1 μm depending on the number of fiducial markers used in the correlation. Arnold *et al*. estimated a success rate of 60 % when milling site-specific lamellas of 200 to 300 nm thick which would drop when aiming for a thickness of 100 to 200 nm.

More recently, ***Gorelick et al. (2019)*** equipped a FIB-SEM with an *in-situ* FM, simplifying sample handling and reducing the risk of sample contamination by limiting the number of cryo-transfer steps. Switching between FM and FIB-SEM imaging modalities required repositioning of the sample inside the vacuum chamber. The 2D positional error made in milling based on FM data is mostly determined by the relocation precision of the sample stage and was reported to be in the range of <420 nm along *X* and <220 nm along *Y*. These relatively large relocation errors prevented high enough localization accuracy to mill a lamella required for high-resolution tomograms, in a targeted fashion.

Coincident imaging with all three microscopes could mitigate the need for fiducial markers and the occurence of relocation errors, thus fully harnessing the power of cryo-CLEM for the cryo-ET workflow. Such a geometry would also allow FM imaging while milling, allowing real-time correction of the milling process in the case of errors, drift, or other misalignments.

Here, we present a coincident 3-beam cryo-CLEM solution, by mounting a tilted light objective lens inside the vacuum chamber such that widefield FM imaging can be done whilst FIB milling. The light microscope is combined with a compact cryogenic microcooler and a custom positioning stage. Our system can be integrated with existing dual beam systems to allow simultaneous, co-incident imaging with both FM, SEM and FIB for *in-situ* FM-guided fabrication of frozen-hydrated lamellae.

## System description

### System Overview

A principal design element of our microscope is the placement of the objective lens (OL) of the fluorescence microscope directly below the sample ***Zonnevylle et al. (2013)***. This minimizes interference with the electron microscope (EM) hardware residing in the upper half space of the EM and offers the following advantages (***Figure 1***): (i) the OL focal point can be aligned to the FIB-SEM working point, allowing simultaneous and coincident imaging with electron-, ion- and photon beams. (ii) *in-situ* FM-guided FIB milling of frozen hydrated lamella without any stage moves can be conducted. (iii) Quality control of the lamella fabrication process can be performed with the light microscope (LM) at every step in the milling process. (iv) Multiple ROIs on the sample can be processed, by centring and focusing the fluorescent ROI in the LM image after a single, initial OL alignment. (v) An OL with a relatively high numerical aperture (NA) can be used, in our case an NA of 0.85, allowing high resolution fluorescence localization.

**Figure 1.**
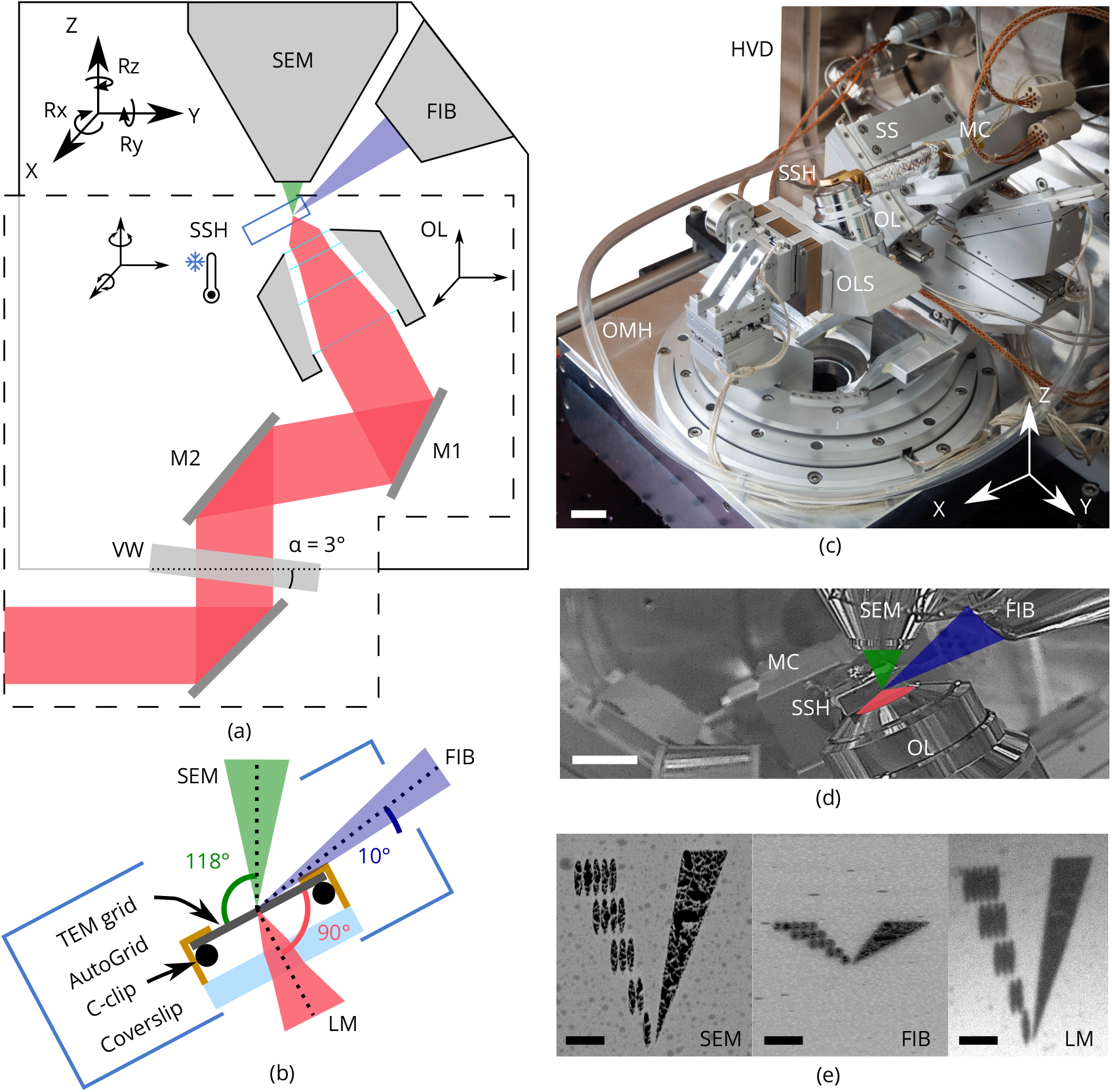
Cryogenic, coincident fluorescence, electron and ion beam microscope. **(a)** Schematic illustration showing the retrofitted hardware (dashed box) separated from the EM hardware in respectively the lower and upper half-spaces of the FIB-SEM. **(b)** Schematic illustration of the sample shuttle holder (SSH) with the TEM grid (sample) clipped into an AutoGrid placed on top of the coverslip. The coincident point is formed by the electron-, ion- and light beams. **(c)** Photograph of the setup showing the HV door (HVD), optical module housing (OMH) on which the sample positioning stage (SS), objective lens stage (OLS), objective lens (OL) and microcooler (MC) are mounted. Scale bar 2 cm. **(d)** Infrared photograph acquired with the charge-coupled device (CCD) camera mounted on the FIB-SEM host system showing the microcooler and objective lens mounted inside the HV chamber. Scale bar 10 mm. **(e)** Images of an alignment pattern in the three imaging modalities, the light microscopy image is acquired by collecting reflected light from the sample. The alignment pattern is milled in the TEM grid coating using the FIB. Scale bar 3.5 μm.

Vitrified cellular samples on TEM AutoGrids ***Rigort et al. (2012)*** are positioned in between the objective lenses of all three microscopes. The AutoGrids are located on top of a thin indium tin oxide (ITO)-coated coverglass that serves as a beam stop to prevent exposure of the OL to electrons or ions. A cryogenic microcooler is used to keep the sample vitrified without the need to cool any of the microscope components, thus retaining unmodified imaging modalities of all three beams. The FM objective lens and microcooler can be moved about in the vacuum by custom piezo positioning stages. A sample shuttle transfer is implemented via a modified PP3006 transfer solution from Quorum Technologies ***Quorum Technologies Ltd (2021)***. The optical microscope hardware resides on a high vacuum (HV) door replacing the original door of the microscope, and fits on a Thermo Fisher Scientific (TFS) SDB chamber, but would also fit FIB-SEM systems from other manufacterers by adapting the design of the door. The piezo stages, microcooler, optical microscope and interface to the host system are controlled using the open-source acquisition software Odemis (Delmic B.V.) ***Piel et al. (2022)***. In the following sections we will discuss the optical microscope, cryogenic sample holder and positioning stage in more detail.

### Optical microscope

To allow optical inspection when the sample is in position for FIB milling at 10° angle of incidence, the OL is in a tilted position to retain perpendicular incidence to the sample surface (***Figure 2)***. It resides in the HV chamber, along with the folding mirrors. All other optical components are mounted inside the optical module at ambient conditions and are separated by an optical vacuum window.

**Figure 2.**
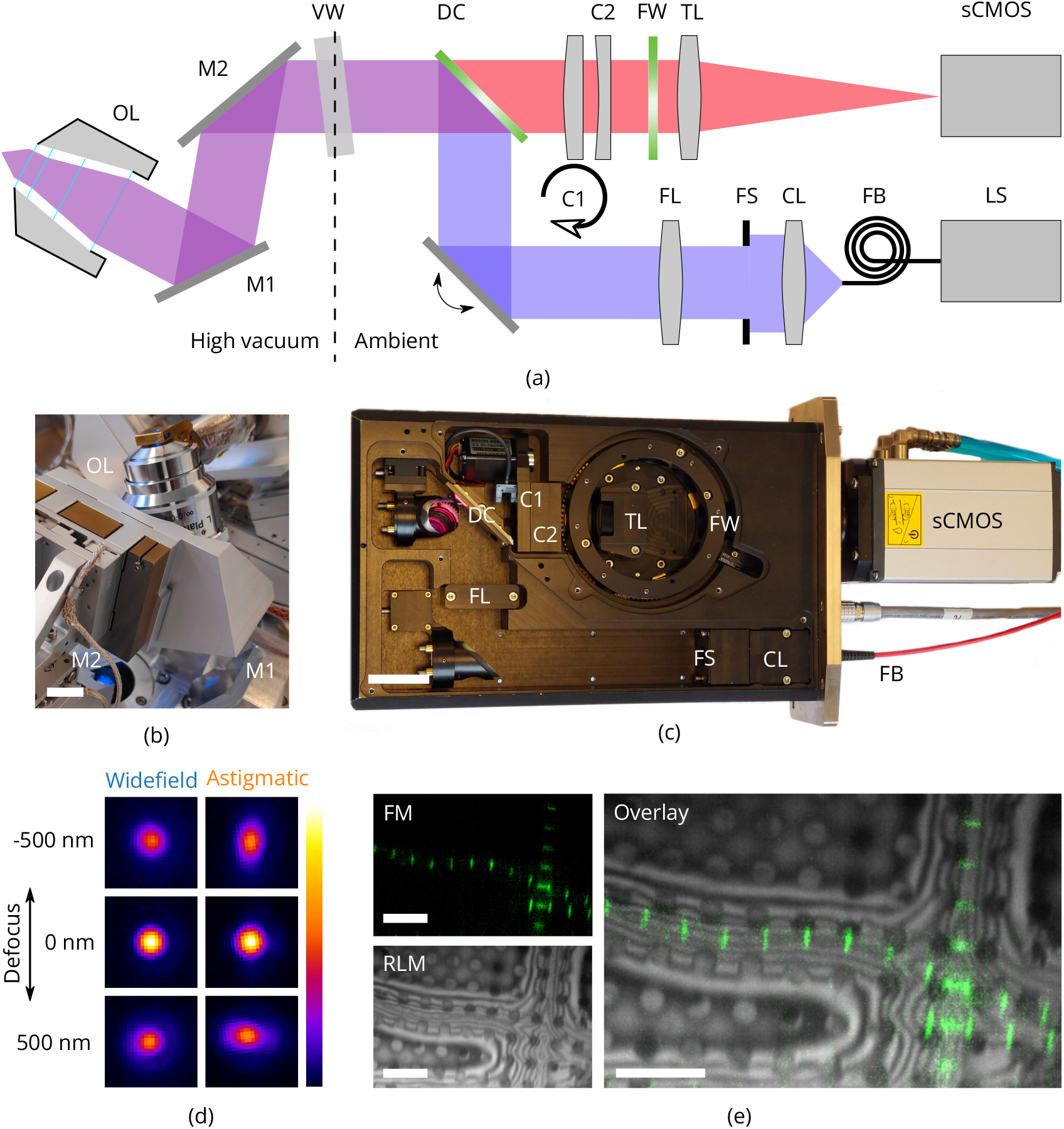
High-resolution fluorescence and reflected light microscopy allows to overlay fluorescence markers on sample structural layout. **(a)** schematic illustration showing the layout of the epi-fluorescence microscope. *M*1 and *M*2 form a folding mirror residing in high vacuum together with the objective lens (OL). The vacuum window (VW) separates the high vacuum from ambient conditions. Excitation and emission paths are separated by the dichroic mirror (DC). Two cylindrical lenses (*C*1 and *C*2) are placed in the emission path to allow for astigmatic point spread function shaping. Emission filters are mounted in the filter wheel (FW), and the tube lens (TL) focuses the image on the camera (sCMOS). The light source (LS) is coupled by a fiber (FB) to the optical module. Light emitted from the fiber is collected by the collector lens (CL), clipped by the field stop (FS) and imaged into the OL object plane by the field lens (FS). **(b)** Optical objective lens mounted on 3DOF stage along with *M*1 and *M*2. Scale bar 1 cm. **(c)** Annotated photograph showing the optical module. Scale bar 4 cm. **(d)** *XY* slices from point spread functions of the optical microscope, at different levels of defocus, both in case of wide field imaging and with induced astigmatic shaping. The point spread functions have been obtained by imaging and averaging individual fluorescent beads (*T*_*sh*_ = 300 K, *N* = 20, λ = 520 nm). Scale bars 1 μm. **(e)** Imaging modalities of the optical microscope. Left top: fluorescence microscope (FM) image of drosophila myofibrils where Sls(700) is tagged with Alexa Fluor 488. Left bottom: reflected light microcopy (RLM) image of the same area as the FM image. Right: Overlay of the FM and RLM images. The circular pattern originates from the Quantifoil R 2/2 film and interference fringes are visible along the radial direction of the myofibrils due to the changing thickness. Scale bars 8 μm.

As the sample is held at cryogenic temperature, a relatively long working distance (WD) objective lens is required. We selected a Nikon CFI L Plan EPI 100xCRA objective lens (Nikon #MUE35900, 100×) for its intermittent NA of 0.85 and working distance ranging between 0.85 to 1.2 mm depending on correction collar setting (0 to 0.7 mm). In our specific case (coverslip #1.5H, 0.170(5) mm thick) the WD is approximately 1 mm. Residual gas analysis with the OL in a separate vacuum setup did not show any foreign species due to the addition of the OL and we did not observe the imaging performance to deteriorate after repetitively pumping and flushing.

In addition to four fluorescence channels, the setup also allows for imaging with RLM by collecting light reflected by the sample from either of the excitation channels through an additional pass-through hole in the filter wheel. Overlaying the FM and RLM images facilitates targeted milling by combining fluorescence signal with contextual information ***Figure 2e***.

Astigmatic point spread function (PSF) shaping is implemented by inserting two obliquely crossed cylindrical lenses, in the form of a Stokes variable power cross-cylinder lens set, in the detection path ***Stokes (1849)***; ***Thompson (1899)***. One of the two lenses is mounted in a rotational mount, actuated by a stepper motor, and is used to switch between astigmatic and regular wide field imaging.

To validate the performance of the optical microscope, we recorded point spread functions at cryogenic conditions by imaging and averaging fluorescent beads (sample temperature 300 K, *N* = 20, λ = 520 nm). The full width half maximum (FWHM) of the widefield PSF is 370 and 1000 nm for the *XY* and *Z* directions respectively. The astigmatic PSF is recorded by setting *θ* = 92°, which leads to an astigmatic deformation in the PSF, ***Figure 2d***.

### Cryogenic sample holder

Conventional cryogenic FIB-SEM systems often use a cold-gas cooled sample holder consisting of a solid block of copper with positioning stage mechanics below it, which is incompatible with our design for the optical microscope. Our cryogenic sample holder (***Figure 3)*** needs to: (i) keep the sample vitrified, (ii) allow the sample to be imaged from three sides simultaneously by SEM, FIB and LM, (iii) accept TEM grids clipped in AutoGrid cartridges, (iv) keep drift and vibration levels at a minimum and (v) shield the sample from contamination as much as possible. We opted for a coldfinger design where the central feature is a customized Joule-Thomson (JT) cryogenic microcooler optimized for its low vibrations, drift and small footprint (DEMCON kryoz), ***DEMCON-kryoz (2021)***; ***Lerou et al. (2008)***. The cooling mechanism is based on high pressure gas undergoing JT expansion through a restriction, producing a maximum net cooling power of ∼200 mW. The cold stage consists of three layers of patterned borosilicate glass (D263T) housed in a titanium tube, and is braided to a cold finger. The 0.1 mm thick titanium tube provides adequate thermal isolation from the bigger aluminium body containing incoming gas lines, electrical wiring and a vacuum connection. The housing is leak tight and differentially pumped by the FIB-SEM vacuum system through a fluoropolymer tube.

**Figure 3.**
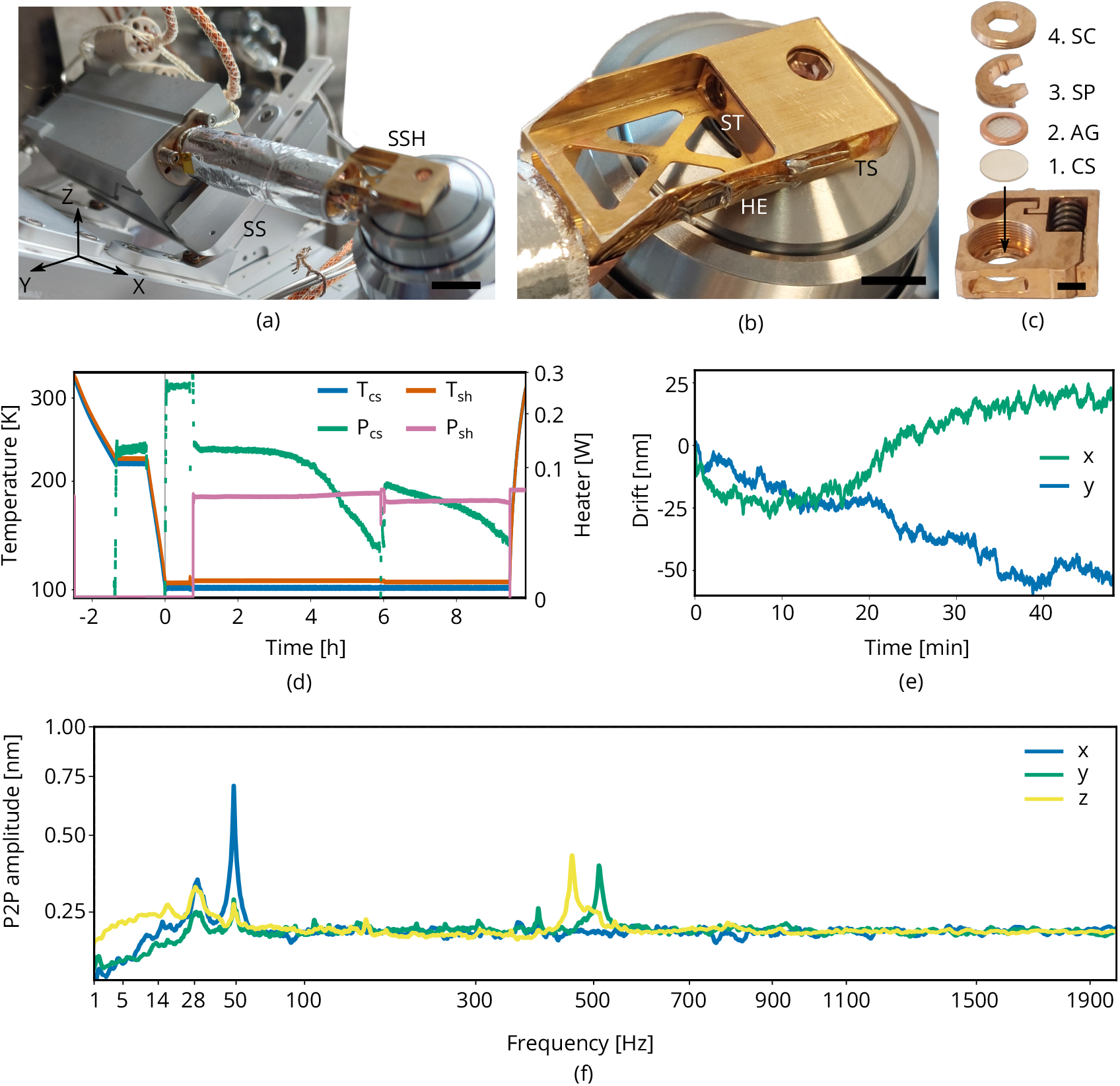
Cryogenic microcooler design and performance. Photographs of **(a)** the microcooler mounted on the sample positioning stage (SS), **(b)** the sample shuttle holder (SSH) in detail and **(c)** the shuttle (ST) and the stacking order of the coverslip (CS), AutoGrid (AG), spacer ring (SP) and screw (SC). Scale bars 10 mm, 5 mm and 2 mm respectively. **(d)** Microcooler thermal performance over time. *T*_*cs*_ and *T*_*sh*_ refer to the cold stage and shuttle holder temperatures (101.5 and 106 K respectively when stabilized). The heater power required for temperature stabilization is also plotted (*P*_*cs*_ and *P*_*sh*_). **(e)** Drift measured whilst regulating both cooler- and shuttle holder temperatures. **(f)** Peak to peak vibration amplitude as a function of frequency at 101 K cooler temperature (*T*_*cs*_) and different directions. These spectra were measured with the microcooler mounted on an aluminium dummy stage inside the FIB-SEM system. The peak at 50 Hz originates from electrical interference.

This compact cooling solution provides local cooling to the sample. We found that it requires approximately two hours to reach <108 K and the cooler can keep the sample cold for more than 9 h. The standing time is solely limited by cryopumping water from residual gas onto the cold surfaces as seen by the decrease in heater power (*P*_*cs*_ and *P*_*sh*_) required for temperature stabilization. The lack of any liquid boil-off keeps the vibration- and drift levels very low. With the shuttle holder temperature stabilized, the drift levels are approximately 1 nm/min along the *y* direction and 0.5 nm/min for the *x* direction. The peak to peak vibration levels are well below 1 nm in all directions.

### Piezo positioning stages

To position the sample within the vacuum chamber, the cryogenic microcooler is mounted on an in-vacuum piezo positioning stage having 5 degrees of freedom: *X, Y, Z, R*_*x*_ and *R*_*z*_. It is built up from individual linear crossed roller bearing positioners using a piezo stick-slip mechanism (SLC Series, SmarAct GmbH) (***Figure 4***) ***SmarAct GmbH (2021)***. By setting *R*_*x*_ = −28°, the FIB maintains a 10° grazing incident angle to the sample surface. The stage has a clear aperture for the optical path and OL with the corresponding piezo positioning stage having 3 degrees of freedom: *X, Y, Z*. The *X* and *R*_*z*_ axis of the sample stage operate independently whilst the *Y, Z* and *R*_*x*_ axis are coupled through a kinematic mount. The *Y* positioners are coupled to three positioners mounted on wedges, rotating them 45° in *Y Z* plane. This geometry was optimized for its stiffness as rotations in *R*_*x*_ require the microcooler and connected tubing to be moved about. *Z* translation is obtained by moving the slanted positioners down whilst compensating the resulting *Y* shift with the *Y* positioners. In the detailed photograph on the right the microcooler mounting plate is shown. It has three ceramic balls on the bottom which mount into three kinematic mounts; two v-grooves and one point contact. Leaf springs on top keep the ceramic balls in place. The stage electronics are programmed such that all sample stage moves are performed in the (possibly rotated) sample coordinate system.

**Figure 4.**
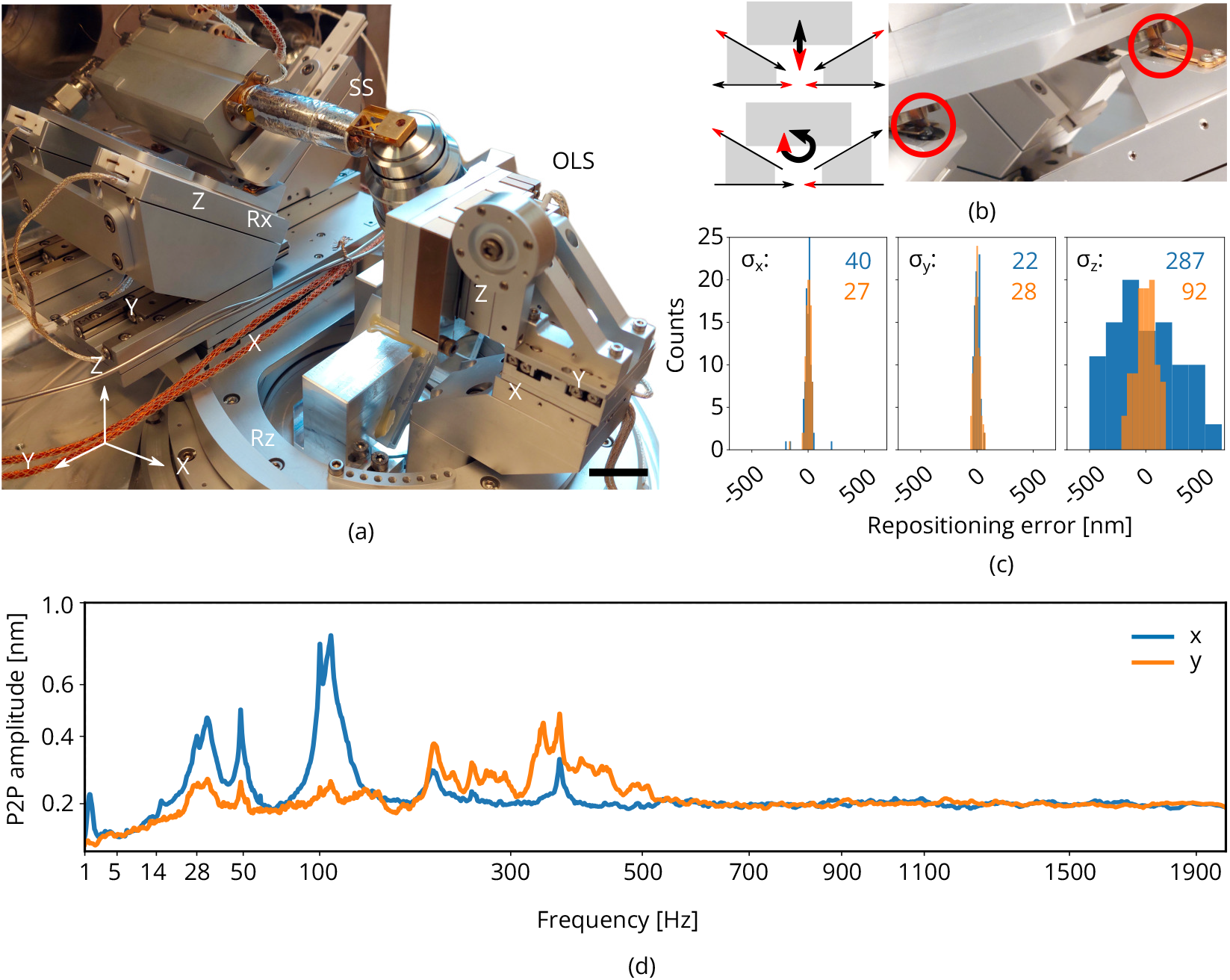
Piezo positioning stage design and performance. **(a)** Photograph of the piezo positioning stages used in the setup. The 5degrees of freedom (DOF) sample stage (SS) moves the microcooler including sample holder around and the 3DOF stage (OLS) positions the OL. Scale bar 2 cm. **(b)** Schematic showing kinematic geometry for movements along *Z* and *R*_*x*_. The microcooler is mounted on an interface plate having three ceramic ball contacts and mount to three kinematic mounts; two v-grooves and one point contact. **(c)** Repositioning error as measured when using the LM to repetitively centering and focusing the same feature in RLM. Images of the sample are acquired with the SEM and FIB and the image to image shifts are used to determine the repositioning error in the *X, Y* and *Z* directions (sample coordinate system). *σ*_*X,Y, Z*_ indicated in blue and orange in nanometer. Number of measurements *N* = 100. **(d)** Peak to peak vibration amplitude as a function of frequency with the microcooler mounted on the 5DOF stage inside the FIB-SEM system. The peak at 50 Hz originates from electrical interference. Due to reduced stiffness of the sample stage (as compared to the aluminium dummy stage), an overall increase is measured whilst staying well below 1 nm peak-to-peak.

We measured the repositioning error of the stage by initiating a random move of the sample positioning stage and then returning to the start position. The same feature was brought into focus with FM (manually, by eye), and centered in the field of view (FOV). We then acquired images of the sample with the SEM and FIB. This procedure was repeated a number of times and the image to image shifts were used to determine the relative error in the *X, Y* and *Z* directions (sample coordinate system). From these measurements we estimated *σ*_*X*_, *σ*_*Y*_ and *σ*_*Z*_ to be 40, 22 and 287 nm respectively (blue) when focusing using a widefield PSF. When using an astigmatic PSF (∼110 mλ, orange) the repositioning error along *Z* is reduced significantly to 92 nm. To probe robustness to mechanical vibrations, we recorded the vibration spectrum at room temperature With the microcooler mounted on the sample stage. The reduced stiffness of the sample stage alters the vibration spectrum, showing more and broader peaks above 100 Hz with respect to the aluminium dummy stage. However, the peak-to-peak vibration levels are well below 1 nm.

## Light microscopy-assisted lamella fabrication in vitrified cells

### Sample loading and 3-beam alignment

With the microcooler at its operating temperature (<108 K), it is moved into the loading position by the piezo stage. Plunge frozen vitrified cells on a clipped TEM grid are loaded in the sample shuttle. The sample shuttle is picked up and transferred in using the modified Quorum transfer system. The sample stage is then positioned such that the TEM grid is brought into the SEM and FIB focus by positioning it near the coincident point of the FIB-SEM. Next, the OL stage is used to bring the sample into the optical microscope focus and care is taken to align the three beams by imaging a piece of bare grid foil with all imaging modalities. The thin (15 to 50 nm) grid foil allows precise alignment of all three beams in the following way: first the sample *Z* height is adjusted such that SEM and FIB image the same area on the sample. Next, the OL position is adjusted such that this area is also imaged in RLM and hence coincidence is achieved with the three beams.

#### Box 1.

**Live LM imaging whilst milling**

**Video 1.**
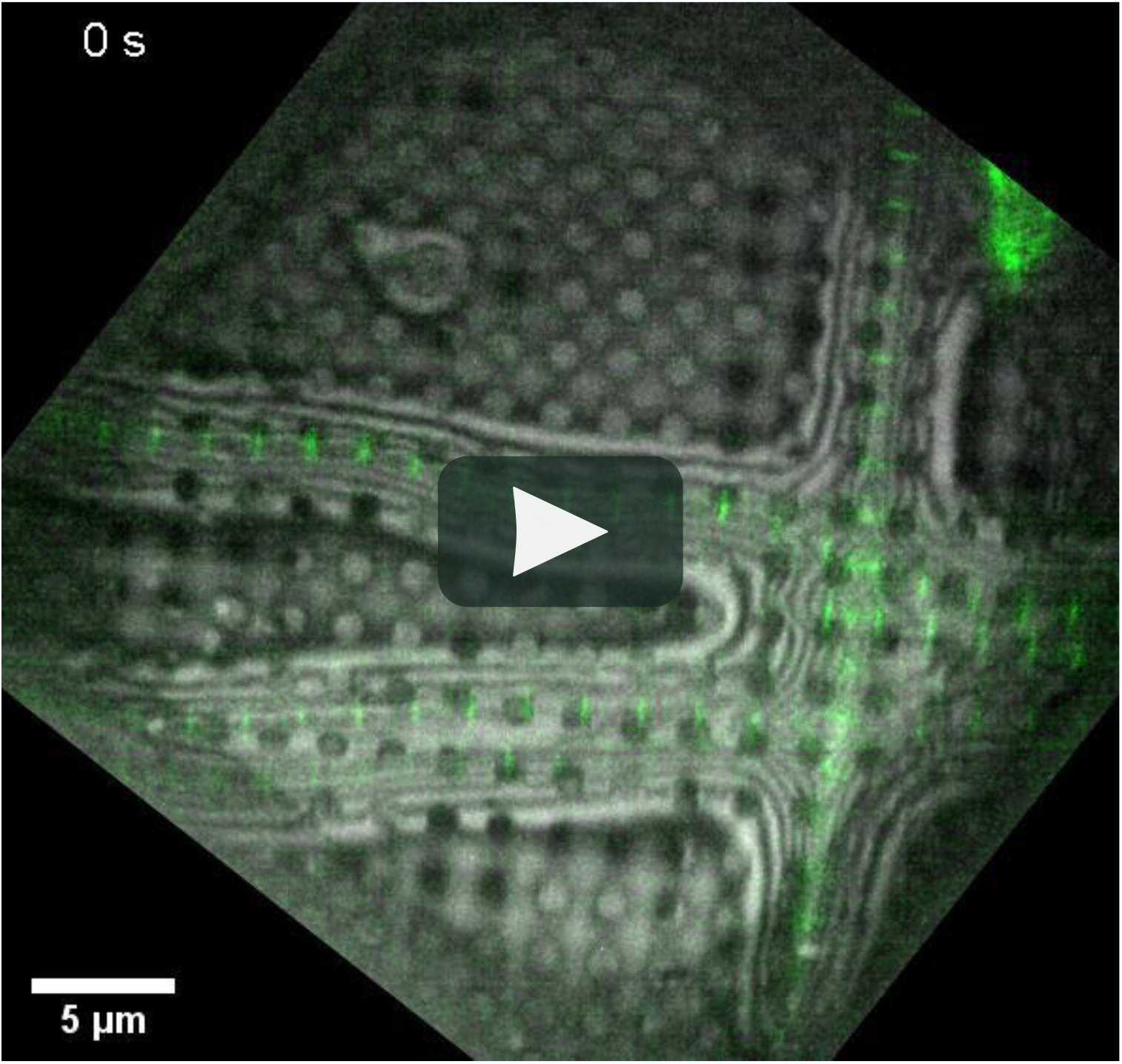
Movie showing RLM and FM images recorded whilst FIB milling a frozen hydrated lamella.

The 3-beam coincident geometry allows FM and RLM images to be acquired during lamella milling in a drosophila myofibril sample. The fluorescence from Sls(700) tagged with Alexa Fluor 488 (green) is overlaid on the reflected light (grayscale).

After this, the objective lens stage is not moved anymore and sample navigation is done solely with the sample positioning stage (*XY*). Bringing a region of interest into the optical microscope focus is also done with the sample stage (*Z*) as *X, Y, Z* moves are performed in the rotated coordinate system of the sample. Having a region of interest in focus and centered in the FOV of the FM/RLM automatically aligns it to the center of the FIB and hence milling can commence.

### Live monitoring and adjustment of FIB milling

FM and RLM images acquired before and during the milling process were used to monitor FIB milling (***Figure 5***). Drosophila and zebrafish myofibrils samples were used for these experiments (Sls(700) tagged with Alexa Fluor 488). The fluorescence (green) is overlaid on the reflected light (grayscale). A few cropped snapshots from this live recording are shown in ***Figure 5a***. The image on the left shows the milling area by the dark blue rectangle and the fluorescent target is red encircled. Four different snapshots of this milling area are shown on the right. The varying color scale is used to indicate the progress in time ranging from 0 to 6 min. In ***Figure 5b***, the intensity of both the FM and RLM signals within the milling area are integrated along *X* and normalized. The dark blue curves are the integrated pixel intensity prior to milling. The light blue peak on the right in the reflection plot corresponds to the second (bottom) myofibril visible in ***Figure 5a***. As the lamella gets thinner, the central reflected intensity peak becomes narrower (going from yellow to red). As the fluorescent target is larger than the lamella thickness, some loss of fluorescence is expected as is shown in the bottom plot. Although the fluorescence signal decreases it still remains after milling. The full video can be found in the Supplementary information.

**Figure 5.**
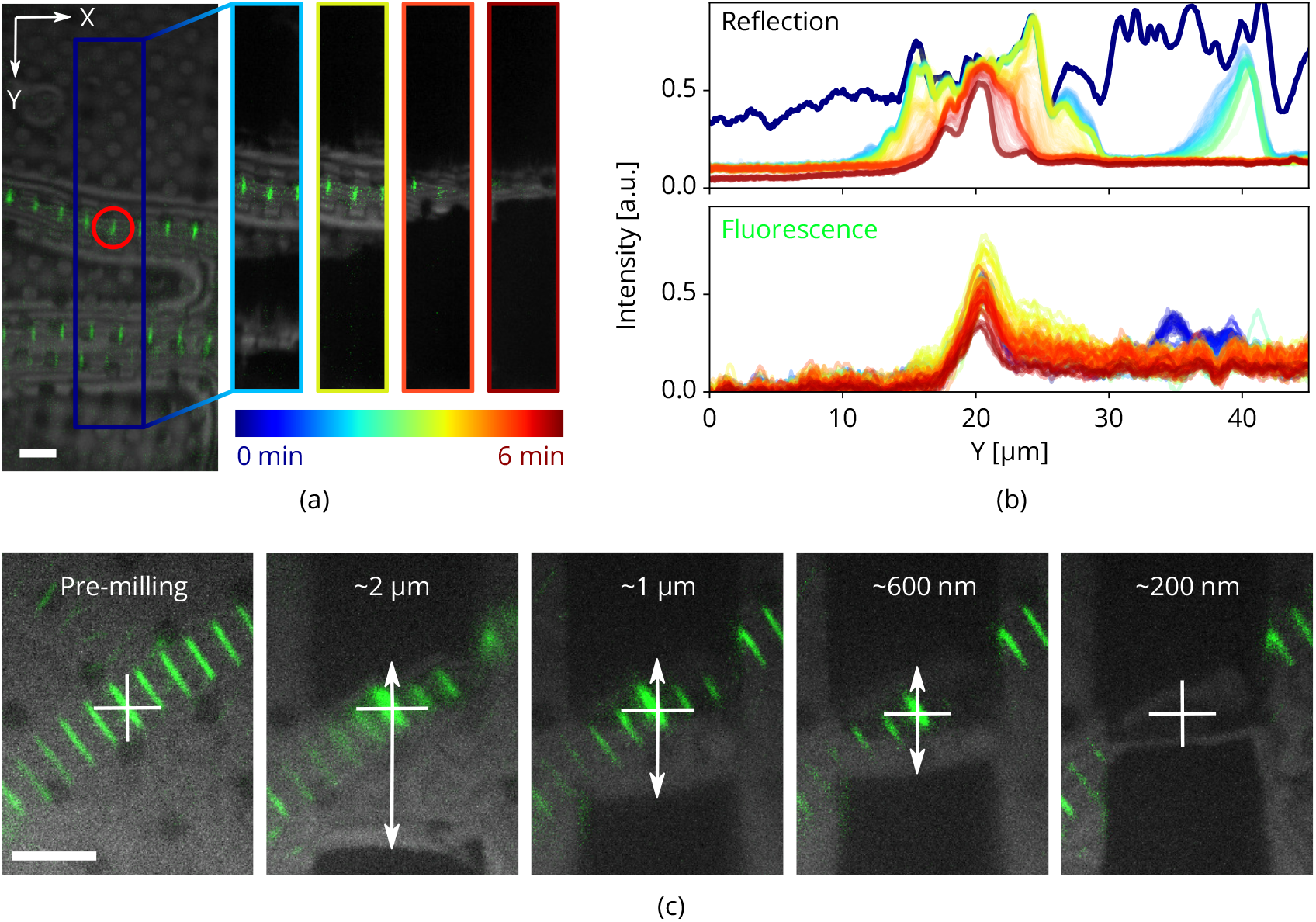
Light assisted FIB milling of frozen hydrated lamellae. **(a)** RLM and FM images were recorded whilst milling. The image on the left shows the indicated milling area marked in dark blue. The four images to the right show snapshots at different stages of milling, denoted with the varying color scale. This color scale is also used in **(b)** where the normalized integrated pixel intensity across the milling area *Y* direction is shown for both the reflection and fluorescence channels. **(c)** Adjustments to the fabrication procedure are easily made when making a lamella based on predefined milling patterns of i. e. 2, 1, 0.6 and 0.2 μm. All scale bars are 5 μm. Drosophila and zebrafish myofibrils in **(a)** and **(c)** respectively. Fluorescence marks the Z-disc of the sarcomere.

Quality control can be performed during every step of the milling procedure without the need for stage movements when switching between the different imaging modalities. Consequently, any misalignment of a feature of interest with respect to the center of the lamella is spotted and corrected for. After each step RLM and FM images are acquired. With the feature of interest marked by the white cross, it is easy to spot a misalignment (white arrows) and correct for this by shifting the milling patterns accordingly. This prevents accidentally milling away the feature of interest and increases sample preparation yield.

### Fluorescence targeted milling and cryo-ET

We illustrate a fluorescence-targeted lamella fabrication workflow by targeting autophagic compartments in HeLa cells (***Figure 6***). We used a red fluorescent protein (RFP)-GFP tandem fluorescenttagged LC3 (mRFP-GFP-LC3) single molecule-based probe that can monitor the autophagosome maturation process ***Kimura et al. (2007)***. This probe emits yellow signals (GFP plus RFP) in the cytosol and on autophagosomes but only red signals in autolysosomes because GFP is more easily quenched and/or degraded in the lysosome than RFP (***Katayama et al. (2008)***). We selected regions of LC3-positive punctae using the mRFP-GFP fluorescence signal and focused this region in the center of the field of view. We then cut the lamella step-by-step and finally polished whilst making sure the feature of interest is still present (***Figure 6a*** through d) at all stages. The contrast was increased to make the fluorescence visible (***Figure 6d***, left) and the final lamella imaged by FIB in the right panel. The sample containing the milled lamella was removed from the FIB-SEM using the transfer system and loaded into the autoloader of the cryo TEM. An overview image of the lamella was acquired and using the RLM image data we overlaid the FM signal guided by the the outline of the lamella as imaged by TEM and RLM. We then acquired tilt series at the sites still showing fluorescence and a z stack is reconstructed the tomograms. No signs of devitrification were visible in these reconstructions, indicating the sample has been well preserved during the entire procedure.

**Figure 6.**
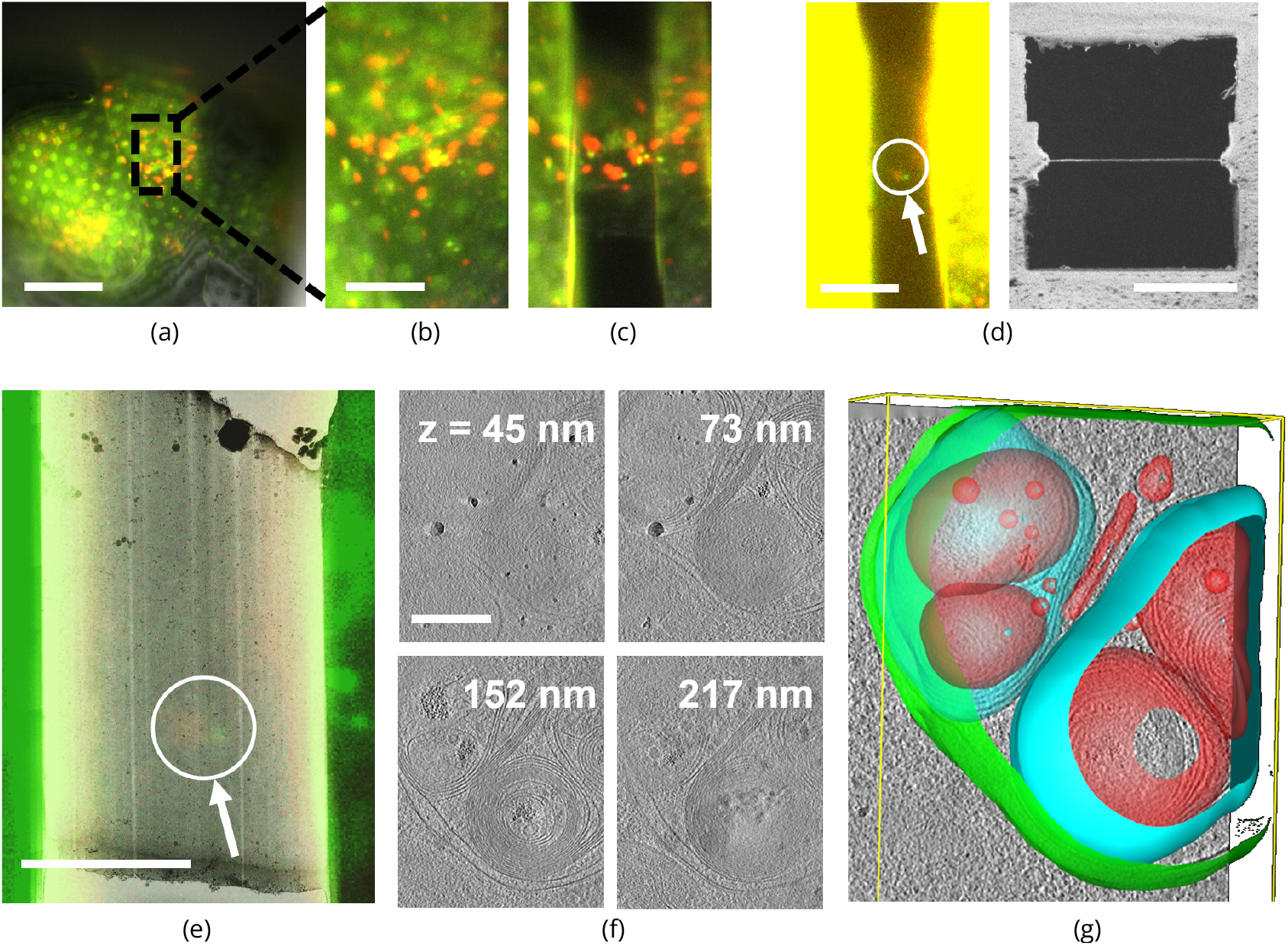
Light targeted FIB milling of frozen hydrated lamellae. **(a)** first the sample is brought into the center of the FOV and focussed properly. Green & yellow: autophagosomes and red: autophagolysosomes. **(b)** Close-up of **(a)** showing the milling area in more detail. **(c)** Snapshot taken at an intermediate milling step. **(d)** left; the lamella finished and polished, imaged in the FM and the contrast enhanced to show the feature of interest. Right; a snapshot acquired with the FIB showing the polished lamella. **(e)** the RLM channel is used to align the light microscope data with the TEM overview image. The fluorescent feature of interest indicates where the tomogram should be acquired. **(f)** *Z* slices of the reconstructed TEM tomogram with **(g)** segmentated membranes. Annotated are autophagolysosome outer membrane (green), autophagolysosome membraneous content (blue and red). Scale bars 40, 10, 10, 5, 5 and 0.25 μm for respectively **(a), (b), (d)** left, **(d)** right, **(e)** and **(f)**.

## Conclusion

We have developed a coincident 3 beam microscope for the cryo-ET workflow which allows direct light microscopy targeted lamella fabrication, without the need for repositioning or fiducial markers. The fluorescence signal intensity from the lamella can be monitored whilst milling making sure that the target remains intact. Any misalignment can be directly observed and corrected. The 5-channel inverted epifluorescent light microscope is diffraction limited in wide field imaging with an NA up to 0.85. Astigmatic point spread function shaping is achieved through the use of a Stokes variable power cross-cylinder lens set. This allows positioning of the sample for targeted milling with errors as small as 25 nm in *XY* and 90 nm in *Z*. The system utilises a novel, liquid nitrogen free cooler design having low levels of vibrations, drift, and an up time of more than nine hours. The entire cryogenic fluorescence microscope can be integrated into a regular FIB-SEM, effectively converting it into a cryo-LM-FIB-SEM, making it possible to add *in-situ* fluorescence targeted milling in an established cryo-ET workflow.

## Outlook

Our approach allows for a live, reflection and/or fluorescence image feed while milling takes place. This opens the additional possibility of an automated milling procedure in which the fluorescent feature of interest is with certainty inside the lamella, before the sample is transferred to the TEM.

We have also implemented a system of two rotatable cylindrical lenses to allow fluorescence imaging with an astigmatic PSF. We have demonstrated that astigmatic imaging improves relocalization accuracy after stage movements to within 90 nm along the optical axis. Further work could push this concept further towards quantitative localization of multiple or potentially even single fluorophores and consequently targeted milling with even lower positional errors. Moreover, an intriguing prospect would be to extend this concept towards cryogenic super resolution microscopy correlated with both SEM and FIB. Potentially, this could also allow for super resolution FM imaging after polishing the lamella, ultimately combining at the single protein level high resolution biological information with the structural data obtained from TEM.

## Material and methods

### Epi-fluorescence microscope

Excitation and emission paths are separated by a dichroic mirror (Semrock #FF410/504/582/669-Di01-25×36), matched by four separate emission filters (Semrock #FF01-440/40-25, #FF01-525/30-25, #FF01-607/36-25, #FF02-684/24-25) in a filter wheel. A 200 mm tube lens (TL) is placed in the center of the filter wheel, and the camera (water cooled Andor Zyla 4.2 PLUS) is fitted on the outside of the optical module. The optical window (Edmund #16-461) separating the HV and ambient environments is mounted at 87° to the optical axis, avoiding a parasitic reflection from excitation light on the camera sensor. A fiber-coupled light emitting diode (LED) excitation light source (Lumencor SPECTRA X central excitation wavelengths: 390, 485, 560 and 648 nm) is used. Light emitted by the fiber is collected by an aspheric collection lens (Edmund #66-312). Together with the field lens (Qioptiq #G322342322), the fiber output is imaged into the exit pupil of the OL. A field stop is placed between the collection and field lenses and is imaged onto the OL object plane, thus restricting the illuminated area on the object (sample). Alignment of the optical path is done by adjusting the 3 DOF kinematic mirror mounts in the excitation and emission paths.

The Stokes variable power cross-cylinder lens set is composed of a plano concave and a plano convex round cylindrical lens (Thorlabs #LK1002RM-A, *f* = −1000 mm & Thorlabs #LJ1516RM-A *f* = 1000 mm). If the the power axes of the two lenses are positioned at right angles (*θ* = 90°), no astigmatism is introduced and hence the PSF remains unmodified. Consequently, variable amounts of astigmatism can be introduced by setting *θ* ≠ 90°.

### Microcooler shuttle holder

The sample shuttle holder mounts to the cold finger protruding at the front of the microcooler. Two temperature sensors (Pt-1000) are present, one residing on the shuttle holder (*T*_*sh*_) and one placed on the micro cold stage (*T*_*cs*_). Near each sensor, a surface-mount technology (SMT) based resistor (68 Ω, International Manufacturing Services #IMS004-3-68) is placed acting as a heater. The thermal gradient between shuttle holder and micro cold stage is ∼5 K, allowing the temperature of both to be regulated semi-independently through control loops. The shuttle contains the AutoGrid (Thermo Fisher Scientific) in which the TEM grid is held in place by a c-clip. Starting with an empty shuttle, first the coverslip (ITO coated D263M glass, 3.4 mm diameter, Schott AG) is placed, followed by the AutoGrid, spacer ring and screw. The shuttle has an actuated leaf spring mechanism which is counteracted when the shuttle is picked up by the transfer rod via a threaded interface. With the transfer rod fully engaged, the leaf spring folds around the main body to reduce the shuttle width so it can move inside the shuttle holder. When unscrewing the transfer rod, the compression spring pushes the leaf spring mechanism outwards, effectively clamping the shuttle inside the holder to ensure good thermal contact between holder and shuttle.

### PSF analysis

Multicolor fluorescent beads (150(30) nm, #FP-0257-2, Spherotech Inc.) were drop casted at low concentration on the glass coverslip. With a multitude of beads (*N* > 20) visible in the field of view, a z-stack was acquired (−3.5 to 3.5 μm, step size of 50 nm, λ = 520 nm, sample temperature 107 K). From this stack the point spread function was measured by averaging multiple bead images using a Python code, as available on GitHub ***Boltje et al. (2022)***. This code works in the following way. The z-stack is loaded as an array and a maximum intensity project along *z* is calculated. Next, TrackPy is used to find all potential features in the 2D image ***Allan et al. (2021)***. The features found are filtered on minimum- and maximum mass to discard noise peaks and clusters of multiple beads such that only single fluorescent beads remain. Beads too close to the edge or to each other are filtered out and the remaining beads are extracted in their respective sub-volumes, based on the NA, the emission wavelength and the pixel size. Any remaining outliers (i.e. doublets or faulty localization) are filtered out by computing the Pearson correlation coefficient between the maximum intensity projections along *z* of all sub-volumes. To localize the remaining beads with sub-pixel precision a 2D Gaussian fit (non-linear least squares) is performed on the sub-volume maximum intensity projection along *z* to get the *x* and *y* position. At this coordinate, a 1D Gaussian fit is performed on the line intensity profile along *z*. Finally, the sub-volumes are up-sampled by a factor of five, the beads are aligned based on the individual *x, y, z* coordinate rounded to the closest up-sampled voxel and are averaged yielding a close approximation of the point spread function of the optical system. To obtain an astigmatic PSF, astigmatism was introduced by setting the cylindrical lens rotation to *θ* = 93.7°, and the procedure for PSF extraction was repeated.

### Vibration measurements

Mechanical vibrations appear when scanning along an edge with clear contrast in the SEM, as is often done to gauge vibration performance ***Jung et al. (2012)***; ***Płuska et al. (2009)***. We use a tin on carbon resolution test sample (#AGS1967T, Agar Scientific Ltd). As the line time determines the sampling frequency we acquire a series of images with different scan resolutions and pixel dwell times, effectively sampling a frequency range of approximately 0.1 to 5000 Hz.

This is done at 0 and 90° scan rotation to measure along the *X* and *Y* directions. Each of the images is analysed line-by-line where a Heaviside step function is used to determine edge position. The Pearson correlation matrix is computed between the original image and a matrix of Heaviside functions having all possible step positions along the horizontal image direction. The vectorized Pearson correlation implementation is used from the GitHub repository belonging to ***Enache et al. (2018)***. The step position versus time can be extracted from the Pearson correlation matrix by taking the absolute value and looking up the indices of the maximum values. The tin ball curvature is removed by applying a Butter high pass filter having a critical frequency *f*_c_ = 5/*T*_*f*_ Hz where *T*_*f*_ is frame time of the image. A scalloping loss corrected FFT is computed, having correct peak-to-peak amplitudes ***Lyons (2011)***. 100+ images are acquired (Helios Nanolab 650 dualbeam, 18 kV, 100 pA beam current, 100 nm horizontal field width, UHR mode) & analysed for each scanning direction, the median of the resulting FFTs is taken to come to the final vibration spectrum. The Python code is available on GitHub ***Boltje and Last (2022)***.

**Figure 7.**
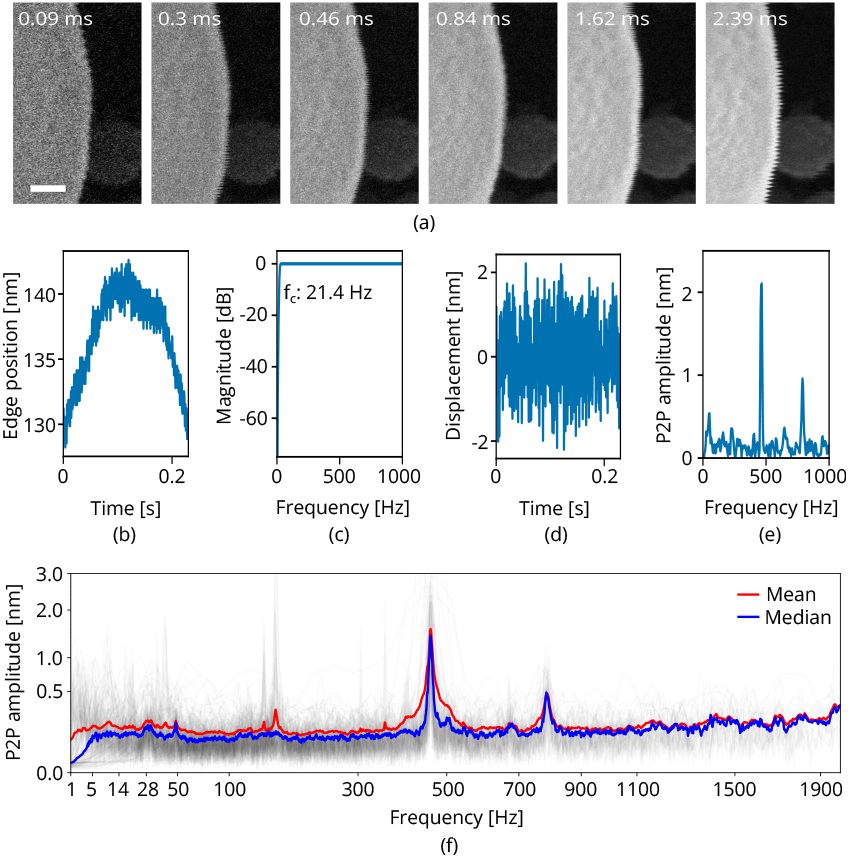
SEM vibration analysis procedure. **(a)** A series of SEM images acquired with different dwell times and thus line times (the latter indicated at the top). **(b)** The edge position as a function of time for one of these images. **(c)** The butter high pass filter through which **(b)** is passed, yielding **(d)** the vibration displacement as a function of time. Finally in **(e)** the scalloping loss corrected fast fourier transform (FFT). Processing all images acquired with the same scan rotation finally yields **(f)** Scale bar 25 nm.

### Repositioning accuracy

The sample stage is positioned such that both SEM and FIB image the metal grid bar on an EM grid. The LM is aligned to image fluorescent beads (150(30) nm, #FP-0257-2, Spherotech Inc.) on the glass cover slip and a single isolated bead is centered in the FOV of the LM. The position is read from the stage encoders and stored. The sample stage is then ordered to make a random 3D move with limits of ±100 and ±30 μm for *XY* and *Z* respectively and moved back to the previous position. The LM is used to manually center and focus the same fluorescent bead in the field of view and after this, images in the FIB and SEM are acquired. This procedure is repeated for a total on 100 times. The image to image shifts are determined by phase cross correlation (as implemented in the Skimage.registration module), for the sequence of FIB and SEM images separately. Due to the viewing geometry (62 and 10° angle of incidence to the sample plane for SEM and FIB respectively) shifts along the image *Y*_i_ direction can be due both sample displacement along *Y*_s_ and *Z*_s_. We extract the sample displacement by minimizing the square error between the two image shifts found (*Y*_SEM_, *Y*_FIB_) and computed image shifts based on the imaging geometry. This yields 99 displacements along the *X, Y* and *Z* sample coordinate system representing the sample stage repositioning accuracy.

### Sample preparation

Drosophila flight muscle myofibrils and zebrafish myofibrils were isolated as described previously by ***Weitkunat and Schnorrer (2014)*** and ***Wang et al. (2021)*** respectively.

HeLa cells stably expressing the mRFP-GFP-LC3 reporter construct (HeLa mRFP-GFP-LC3; kind gift from Fulvio Reggiori, University of Groningen, Netherlands) were cultured in Dulbecco’s Modified Eagle Medium (DMEM), 10 % fetal calf serum, 1 % pen/strep supplemented with 0.6 μg/mL G418 at 37 °C with 5 % CO2 and a humidified atmosphere. Prior to seeding on grids, cells were washed three times with warm medium, followed by two washing steps with PBS. 1 mL of trypsin was added, followed by a 5 min incubation at 37 °C, in a 5 % CO2 humidified atmosphere. Cells were seeded at 660 000 per tissue culture dish (60 mm; Sarstedt). After a brief cleaning with ethanol and pre-washing with DMEM medium, microscopy slides were placed within the culture dish. Glow-discharged carbon coated gold mesh grids (QF 2/2 AU 200, Quantifoil) were placed on the edge of the glass microscopy slides, and cells were gently dispensed into the dish. Cells were cultured for an additional 18 h and then washed twice with pre-warmed PBS to remove any remaining culture medium. Then 5 mL of warm starvation buffer (20 mM HEPES pH 7.5, 140 mM NaCl, 1 mM CaCl2, 1 mM MgCl2, 10 mM glucose) was added. After 2 h at 37 °C, 5 % CO2, the cells were imaged using the EVOS cell imaging system (Thermo Fisher Scientific). Gold mesh grids were removed from the tissue culture dish and immediately prior to blotting, 3 μL of warm starvation buffer was added to the grid. Blotting conditions on the Leica GP2 vitrification robot were blotting from the back for ∼2 s in a chamber with 98 % humidity at 21 °C and then plunged into liquid ethane.

### 3-beam alignment and fluorescence targeted milling in cryo-FIB-SEM

Validation experiments using the system have been carried out at the Department of Structural Biochemistry, Max Planck Institute of Molecular Physiology in Dortmund. The original HV door of the Aquilos Cryo-FIB (Thermo Fisher Scientific) was replaced with our HV door, after which the system was evacuated and and pumped for several days. Prior to experiments, the preparation station in the low-humidity glovebox, the cryogenic chamber shield and the microcooler were cooled down ***Tacke et al. (2021)***. Cellular samples on clipped EM grids were mounted into the shuttle inside the glovebox, picked up with the modified Quorum transfer system and transferred into the system via a load lock. After loading, the sample stage was homed and moved to the coating position, where the gas injection system is used to apply a Pt precurser coating to the EM grid through the top SEM access hole in the shuttle. Afterwards, the sample stage is moved to the predefined 3-beam imaging position, followed by the OL stage.

Due to the large distance traversed, the 3-beam alignment needs to be refined after loading a sample, and is done by imaging a piece of bare grid foil with all imaging modalities. The thin (15 to 50 nm) grid foil allows precise alignment of all three beams: (i) The sample *Z* height is adjusted such that SEM and FIB image the same area on the sample (i.e. eucentric height, the sample is in the coincident point of the SEM and FIB beams). (ii) The OL position is adjusted such that this area is also imaged in RLM and hence coincidence is achieved with the three beams. (iii) A small hole (2 μm diameter) is milled through the grid foil, using the FIB. (iv) This hole is used as an alignment feature, effectively refining the 3-beam coincidence as outline above. This process takes about 5 to 10 min, and after this sample navigation and focusing is done through the sample stage. Having a region of interest in focus and centered in the FOV of the FM/RLM automatically aligns it to the center of the FIB. Sites for milling were identified by first identifying them in a low magnification SEM overview image, followed by a more thorough inspection with the light microscope (both FM and RLM). With a selected region of interest in focus and centered into the LM field of view, FIB milling can commence, starting with the first rough milling step along with the relief cuts.

The SEM is used to assess the lamella thickness, and when thin enough, a final inspection is done with the LM. With 3 to 6 milling sites complete, the sample was transferred out of the system and into the glovebox. The AutoGrid was taken out of the shuttle and placed into the autoloader cassette which was loaded into the NanoCab (Thermo Fisher Scientific) and transferred to the electron cryo microscope.

### Cryo-electron tomography and tomogram reconstruction

To avoid contamination, grids containing milled lamellae were transferred through a low-humidity glovebox ***Tacke et al. (2021)***, into a Titan Krios transmission electron microscope (Thermo Fisher Scientific), equipped with a K3 camera (Gatan) and an energy filter. Projection images were acquired using SerialEM software ***Mastronarde (2005)***. Overview images of were acquired at 4800× or 8700× nominal magnification to identify locations for high-magnification tilt series acquisition. Tilt series were acquired at 42 000 ×, 53 000 × or 81 000 × nominal magnification (pixel size 2.32 Å, 1.81 Å and 1.18 Å respectively). A dose-symmetric tilting scheme ***Hagen et al. (2017)*** was used during acquisition with a tilt range of −56° to 56° relative to the lamella plane at 2° increments. A total dose of 130 to 150 e^−^ /Å^2^ was applied to the sample.

Individual tilt movies acquired from the microscope were motion corrected, contrast transfer function corrected and combined into stacks using Warp ***Tegunov and Cramer (2019)***. The combined stacks were then aligned and reconstructed using IMOD ***Kremer et al. (1996)***.

## Author contributions

SH conceived and initialized the collaboration between the different research groups. Initial ideas and insight into the cryo-ET workflow was provided by JPH, AJJ, GJJ, AJK, JMP, SR and RW. The majority of the conceptual system design was done by DBB with feedback from JPH, AJJ, GJJ, AJK, JMP, SR, RW and SH. DBB, CTHJ and MGFL, planned and carried out the experiments. ST and ZW assisted in doing cryo-FIB milling and performed cryo-ET experiments. EBW helped with light microscopy experiments and PSF analysis. MJK performed the stage reposition measurements. AJJ and CAP prepared the HeLa samples whilst ZW provided the myofibril samples. DBB processed and analyzed the data with input from JPH, AJJ, CTHJ, MGFL, ST, ZW and EBW. DBB wrote the manuscript with input from EBW, AJJ, and JPH. All authors discussed the results and commented on the manuscript.

## Acknowledgments

Drosophila flight muscle myofibrils were kindly provided by E.H. Chan & F. Schnorrer (Institut de Biologie du Développement de Marseille), and zebrafish myofibrils by Y. Hinits & M. Gautel (King’s College London). We express our gratitude to Fulvio Reggiori (University of Groningen, Netherlands) for providing the HeLa cells and are thankful to Mingjun Xu for help during sample preparation. We thank Andries Effting (Delmic B.V.) for helpful discussions, and we would like to thank Ryan Lane (TU Delft) for his contribution in various Python developments. The majority of the 3D CAD design was done by Thomas van der Heijden (Delmic B.V.), for which we are gratefull. This work was financially supported by; NWO-TTW project №17152 to JPH, NIH grant RO1 AI127401 to GJJ, European SME2 grant №879673 to Delmic B.V. and Eurostars grant №E13008 to SH and SR.

## References

Allan DB, Caswell T, Keim NC, van der Wel CM, Verweij RW. soft-matter/trackpy: Trackpy v0.5.0; 2021, https://doi.org/10.5281/zenodo.4682814, doi: 10.5281/zenodo.4682814.

Arnold J, Mahamid J, Lucic V, De Marco A, Fernandez JJ, Laugks T, Mayer T, Hyman AA, Baumeister W, Plitzko JM. Site-specific cryo-focused ion beam sample preparation guided by 3D correlative microscopy. Biophysical journal. 2016; 110(4):860–869.

Boltje DB, Last MFG. Mechanical-Analysis; 2022, https://github.com/hoogenboom-group/Mechanical-Analysis.

Boltje DB, van der Wee EB, Lane R. PSF-Extractor; 2022, https://github.com/hoogenboom-group/PSF-Extractor.

Buckley G, Gervinskas G, Taveneau C, Venugopal H, Whisstock JC, de Marco A. Automated cryo-lamella preparation for high-throughput in-situ structural biology. Journal of structural biology. 2020; 210(2):107488.

Chiang EY, Hidalgo A, Chang J, Frenette PS. Imaging receptor microdomains on leukocyte subsets in live mice. Nature methods. 2007; 4(3):219–222.

DEMCON-kryoz. DEMCON kryoz – Kryoz Technologies; 2021, https://www.demcon-kryoz.nl/ (Retrieved 13-09-2021).

Enache O, Lahr D, Natoli T, Litichevskiy L, Wadden D, Flynn C, Gould J, Asiedu J, Narayan R, Subramanian A. The GCTx format and cmap {Py, R, M} packages: resources for the optimized storage and integrated traversal of dense matrices of data and annotations. bioRxiv.[Google Scholar]. Human Molecular Genetics. 2018; 27(R1).

Gorelick S, Buckley G, Gervinskas G, Johnson TK, Handley A, Caggiano MP, Whisstock JC, Pocock R, de Marco A. PIE-scope, integrated cryo-correlative light and FIB/SEM microscopy. Elife. 2019; 8:e45919.

Hagen WJ, Wan W, Briggs JA. Implementation of a cryo-electron tomography tilt-scheme optimized for high resolution subtomogram averaging. Journal of structural biology. 2017; 197(2):191–198.

Hylton RK, Swulius MT. Challenges and triumphs in cryo-electron tomography. Iscience. 2021; 24(9):102959.

Jung KO, Kim SJ, Kim DH. An approach to reducing the distortion caused by vibration in scanning electron microscope images. Nuclear Instruments and Methods in Physics Research Section A: Accelerators, Spectrometers, Detectors and Associated Equipment. 2012; 676:5–17.

Katayama H, Yamamoto A, Mizushima N, Yoshimori T, Miyawaki A. GFP-like proteins stably accumulate in lysosomes. Cell structure and function. 2008; 33(1):1–12.

Kimura S, Noda T, Yoshimori T. Dissection of the Autophagosome Maturation Process by a Novel Reporter Protein, Tandem Fluorescent-Tagged LC3. Autophagy. 2007; 3(5):452–460. https://doi.org/10.4161/auto.4451, doi: 10.4161/auto.4451, pMID: 17534139.

Koning RI, Koster AJ, Sharp TH. Advances in cryo-electron tomography for biology and medicine. Annals of Anatomy-Anatomischer Anzeiger. 2018; 217:82–96.

Kremer JR, Mastronarde DN, McIntosh JR. Computer visualization of three-dimensional image data using IMOD. Journal of structural biology. 1996; 116(1):71–76.

Lerou PPPM, ter Brake HJM, Jansen HV, Burger JF, Holland HJ, Rogalla H. Micromachined Joule-Thomson Coolers. AIP Conference Proceedings. 2008; 985(1):614–621.

Lyons R. Reducing FFT scalloping loss errors without multiplication. IEEE Signal Processing Magazine. 2011; 28(2):112–116.

Mastronarde DN. Automated electron microscope tomography using robust prediction of specimen movements. Journal of structural biology. 2005; 152(1):36–51.

Piel E, de Laat R, Tsitsikas K, Winkler P, Muskens A, Rossberger S, Pals T, Mavrikopoulou V, Kleijwegt K, Lazem B, Barazi M, Helsloot A. Open Delmic Microscope Software; 2022, https://www.delmic.com/.

Płuska M, Czerwinski A, Ratajczak J, Katcki J, Oskwarek Ł, Rak R. Separation of image-distortion sources and magnetic-field measurement in scanning electron microscope (SEM). Micron. 2009; 40(1):46–50.

Quorum Technologies Ltd. PP3006 CoolLok brochure; 2021, v4.

Rigort A, Bäuerlein FJ, Villa E, Eibauer M, Laugks T, Baumeister W, Plitzko JM. Focused ion beam micromachining of eukaryotic cells for cryoelectron tomography. Proceedings of the National Academy of Sciences. 2012; 109(12):4449–4454.

Schaffer M, Pfeffer S, Mahamid J, Kleindiek S, Laugks T, Albert S, Engel BD, Rummel A, Smith AJ, Baumeister W, et al. A cryo-FIB lift-out technique enables molecular-resolution cryo-ET within native Caenorhabditis elegans tissue. Nature methods. 2019; 16(8):757–762.

SmarAct GmbH. High Precision Positioning and Metrology Solutions - SmarAct; 2021, https://www.smaract.com/index-en (Retrieved 13-09-2021).

Stokes G. On a mode of measuring the astigmatism of a defective eye. Rep Br Assoc. 1849; 2:10–11.

Tacke S, Erdmann P, Wang Z, Klumpe S, Grange M, Plitzko J, Raunser S. A streamlined workflow for automated cryo focused ion beam milling. Journal of Structural Biology. 2021; 213(3):107743.

Tegunov D, Cramer P. Real-time cryo-electron microscopy data preprocessing with Warp. Nature methods. 2019; 16(11):1146–1152.

Tegunov D, Xue L, Dienemann C, Cramer P, Mahamid J. Multi-particle cryo-EM refinement with M visualizes ribosome-antibiotic complex at 3.5 Å in cells. Nature Methods. 2021; 18(2):186–193.

Thompson SP. On Obliquely-crossed Cylindrical Lenses. Proceedings of the Physical Society of London (1874-1925). 1899; 17(1):81.

Turk M, Baumeister W. The promise and the challenges of cryo-electron tomography. FEBS letters. 2020; 594(20):3243–3261.

Villa E, Schaffer M, Plitzko JM, Baumeister W. Opening windows into the cell: focused-ion-beam milling for cryo-electron tomography. Current opinion in structural biology. 2013; 23(5):771–777.

Vulović M, Ravelli RB, van Vliet LJ, Koster AJ, Lazić I, Lücken U, Rullgård H, Öktem O, Rieger B. Image formation modeling in cryo-electron microscopy. Journal of structural biology. 2013; 183(1):19–32.

Wang Z, Grange M, Wagner T, Kho AL, Gautel M, Raunser S. The molecular basis for sarcomere organization in vertebrate skeletal muscle. Cell. 2021; 184(8):2135–2150.

Weitkunat M, Schnorrer F. A guide to study Drosophila muscle biology. Methods. 2014; 68(1):2–14.

Wolff G, Limpens RW, Zheng S, Snijder EJ, Agard DA, Koster AJ, Bárcena M. Mind the gap: Micro-expansion joints drastically decrease the bending of FIB-milled cryo-lamellae. Journal of Structural Biology. 2019; 208(3):107389.

Zimmerli CE, Allegretti M, Rantos V, Goetz SK, Obarska-Kosinska A, Zagoriy I, Halavatyi A, Hummer G, Mahamid J, Kosinski J, et al. Nuclear pores dilate and constrict in cellulo. Science. 2021; 374(6573):eabd9776.

Zonnevylle A, Van Tol R, Liv N, Narvaez A, Effting A, Kruit P, Hoogenboom J. Integration of a high-NA light microscope in a scanning electron microscope. Journal of microscopy. 2013; 252(1):58–70.

